# DiviSSR: Simple arithmetic for efficient identification of tandem repeats

**DOI:** 10.1101/2021.10.05.462997

**Authors:** Akshay Kumar Avvaru, Rakesh K Mishra, Divya Tej Sowpati

## Abstract

Numerical or vector representations of DNA sequences have been applied for identification of specific sequence characteristics and patterns which are not evident in their character (A, C, G, T) representations. These transformations often reveal a mathematical structure to the sequences which can be captured efficiently using established mathematical methods. One such transformation, the 2-bit format, represents each nucleotide using only two bits instead of eight for efficient storage of genomic data. Here we describe a mathematical property that exists in the 2-bit representation of tandemly repeated DNA sequences. Our tool, DiviSSR (pronounced divisor), leverages this property and subsequent arithmetic to achieve ultrafast and accurate identification of tandem repeats. DiviSSR can process the entire human genome in ∼30s, and short sequence reads at a rate of >1 million reads/s on a single CPU thread. Our work also highlights the implications of using simple mathematical properties of DNA sequences for faster algorithms in genomics.

**Availability:** DiviSSR can be installed directly using python pip. The source code and documentation of DiviSSR are available at https://github.com/avvaruakshay/divissr.git.

## Introduction

Many functionally important genomic regions display distinctive sequence characteristics. Identifying these patterns and signatures often becomes the first step towards understanding their function. Numerical or vector transformations of nucleotide sequences introduce and/or reveal a mathematical structure to certain genomic elements. Mathematical models that capture these structures have previously been used in determining the structural, thermodynamic, and bending properties of DNA [1], biological sequence querying [2], estimating DNA sequence similarity [3-5], sequence alignment [6], and identification of repetitive sequences [7]. A relatively recent transformation approach called the 2-bit format represents the 4 nucleotides as a combination of 2 bits. This format has been pervasively used in genomics for efficient storage of DNA sequence information. Here, we leverage a mathematical property introduced by integer representation of nucleotides for identification of tandemly repeated DNA sequences.

Microsatellites, also known as Simple Sequence Repeats (SSRs) or Short Tandem Repeats (STRs) are a class of DNA tandem repeats with a repeating motif size of 1-6bp. SSRs are shown to have non-random distribution in eukaryotic genomes [8], and display a higher rate of polymorphism due to replication errors, particularly replication strand slippage. Abnormal polymorphisms of microsatellites are associated with several human genetic diseases including Huntington’s disease, Friedrich’s ataxia, Spinocerebellar ataxia, Myotonic dystrophy, and many others. Apart from repeats at specific genomic locations, studies point to their broader roles in gene regulation and genome organisation [9-12]. More recently, variations in a subset of STRs called eSTRs have been linked to the expression of proximal genes [13].

Given the ubiquitous roles of SSRs, their efficient identification has been a long standing goal in computational biology. Previous approaches have accomplished this either by sequence matching or using predictive models to measure repetitive properties of the sequences. Here we propose a novel approach defining an arithmetic rule associated with integer transformations of DNA tandem repeats. Our tool, DiviSSR, identifies tandem repeats by applying a division rule on the binary numbers resultant after 2-bit transformations of DNA sequences. DiviSSR is on average 5-10 fold faster than the next best tools and takes just ∼30 secs to identify all perfect microsatellites in the human genome. The performance of the algorithm is motif size agnostic and takes constant time for any motif size(s) on a given genome. The stable version of DiviSSR is publicly available on PyPI, and its source code is deposited at https://github.com/avvaruakshay/divissr.git.

## Materials and Methods

The integer representation of a nucleotide sequence is mapping of each nucleotide to an integer. We map the four nucleotides A, C, G, T to the first four integers 0, 1, 2, 3, a convention also followed by the widely used 2-bit format for efficient storage of DNA sequences. This integer representation of DNA tandem repeats conforms to a unique division rule, as explained in subsequent sections. DiviSSR scans the genome in windows, converts each window sequence to 2-bit format, and checks if the resulting number qualifies the division rule. The detailed explanation of the algorithm is below and an overview of the method is depicted in Figure 1.

**Figure 1:**
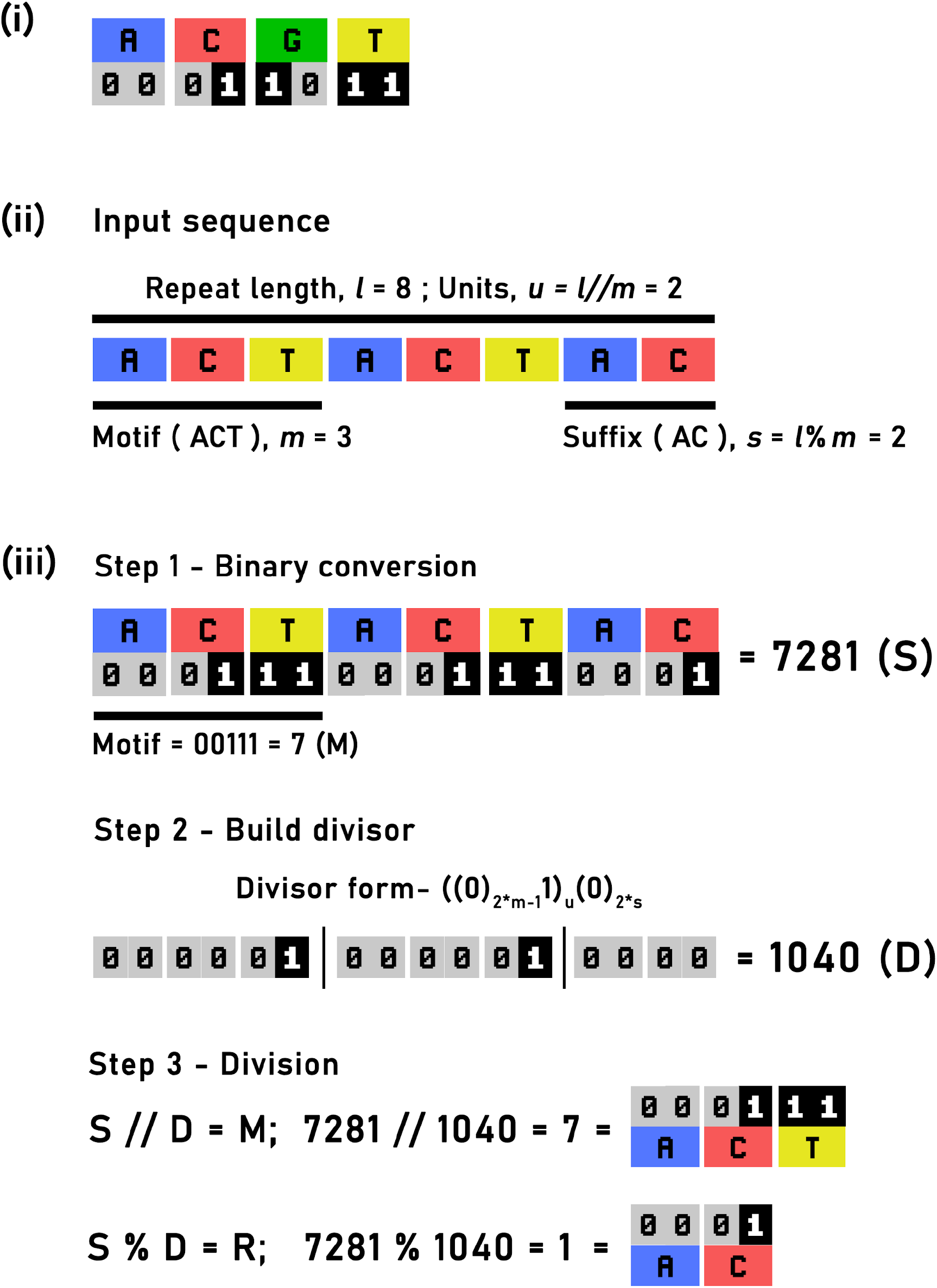
i) 2-bit representation of each nucleotide. ii) Example repeat sequence of ACT motif (*m*=3) with 2 complete units (*u*=2) and a suffix of 2 bases (*s*=2). iii) Binary representation of the repeat sequence generates a number denoted by *S*, which here is 7281. The binary representation of the motif, ACT, yields the number 7 and is denoted by M. The divisor (*D*) is built based on the lengths of the repeat, the motif and the suffix and is of the numerical form ((0)_2*m-1_1)(0)_2*s_. The value of D is 1040 here. The division of S with D yields a quotient, which is 7, equal to binary representation of the motif and the remainder equals binary representation of partial suffix.

### The DiviSSR number

A repeat number, *N*, is a number of *n* digits formed by *u* times tandem repetition of a smaller number *M* of *m* digits, where 1 ≤ *m* ≤ *n*/2. Such repeat numbers can be factored into their motif representations. For example, the repeat number 123123123, with the tandem occurrence of motif 123 thrice, can be represented as (123*10^6^ + 123*10^3^ + 123*10^0^) = 123*(1001001). A similar repetitive number of a 3-digit motif, 865865865, can be factored as 865*(10^6^ + 10^3^ + 10^0^) = 865*(1001001). The number in parentheses, which we call the DiviSSR number, will be the same for all repeat numbers where *n* = 9, *m* = 3, and *u* = 3. Inversely, any 9 digit number perfectly divisible by this DiviSSR is a repeat number of a 3 digit motif repeated thrice. The DiviSSR number is unique for every combination of *u* and *m*, and takes the form (0_*m*-1_1)_*u*_, i.e. *u* repetitions of the motif (*m* - 1) 0s suffixed by a 1. Repeat numbers which end in a partial motif *S*, a number with first *s* digits of *M* where 1 ≤ *s* < *m*, are not perfectly divisible by *M* but leave *S* as the remainder. For example, 86586586586 can be factored as 865*(100100100) + 86. In such cases the divisor will be *u* repetitions of motif 0_*m*-1_1 followed by *s* zeros i.e., (0_*m*-1_1)_*u*_ 0_*s*_. The generalisation for all repeat numbers is *N* = (*M*)_*u*_ *S* = (0_*m* -1_1)_*u*_ 0_*s*_ + *S* (Fig S1). This division property is exhibited by repeat numbers of any number base, including binary numbers. DiviSSR scans the query DNA in sections of overlapping *k*-mers, converts each *k*-mer into corresponding binary number and conducts the division check, as explained below.

### Repeat identification

Given a maximum motif size m”, a repeat length cutoff of k, and *k* ≥ 2\**m’’*, DiviSSR scans the sequence through a window of *k* bp length by shifting 1bp at each step. The sequence in the window is converted into 2-bit format and stored as an integer. For each motif size *m* in the non-redundant set (see below), the divisor is calculated. The divisor for a check of motif size *m* takes the form (0_2\**m*-1_1)0_2\**s*_, as each nucleotide is represented by 2 bits. The converted binary number undergoes a division check once for each motif size among the pre-built non-redundant set in the descending order. If a window fails the check for all motif sizes, the program moves to the next window. If the window passes the check, the motif size and the start position of the window are recorded. Motif size is used to retrieve the sequence of the motif from the binary number. True atomicity of the motif and the repeat class (explained in supplementary) of the motif is checked from its sequence. The program moves to the next window to check for possible continuation of the repeat. Two consecutive windows can pass the division rule check only if both the windows are repeats of the same motif. Two repeat sequences of two different motifs with motif sizes *m1, m2* can at max overlap by (min(*m1, m2*))-1 bp. Consecutive windows overlap by *k*-1bp and *k* is at least twice the maximum motif size. This strictly confirms that consecutive windows passing the checks are not repeat sequences of different motifs. Once the division check fails for a given window, the end position of the repeat is recorded. The repeat is reported and the program moves further in processing the sequence completing the process till the end of the sequence.

### Checks for atomicity and redundancy

The repeat check for a particular motif size (*m*) suffices as a check for all repeats with motif sizes which are factors of *m*. For the user-defined motif size range of *m’* to *m’’* (*m’’* > *m’*) DiviSSR checks for factor-multiple relationships between all pairs of numbers within the range. A non-redundant set of motif sizes is built by dropping the smaller factors. At the time of identification, the true atomicity of the repeat is lost because of the reduction in the checks. For each identified repeat DiviSSR identifies the true atomicity of the repeat by checking if the identified motif can be further factored as the repeat of smaller motif size.

### Downstream analysis

DiviSSR is equipped with two auxiliary modules for downstream analysis of repeats. One module annotates repeats in genomes for which feature information is provided in GTF/GFF formats. The annotation is done with respect to the closest gene from each repeat. This is implemented using an in-house Python script. A descriptive explanation of the annotation algorithm is explained in Supplementary Methods. The second module builds an interactive HTML report summarising the trends of repeat distribution as interactive tables and plots. A static native HTML template is used for the report and the data visualisations are generated using echarts (https://echarts.apache.org/en/index.html), a JavaScript plotting library. The HTML report is input file-format aware (i.e. genome or raw sequence dataset) and displays a slightly different output for both.

### Comparison with other tools

We have compared DiviSSR’s performance with other perfect repeat identification tools, Kmer-SSR[15], Mreps[14], PERF[16], SSRIT[17], and Misa[18]. The comparison was limited to the tools which were based on deterministic and exhaustive algorithms. We employ two tests for comparison of run time profiles based on the input genome size and the desired motif length ranges across tools. All tests were run on a machine with Intel® Core™ i7-4810MQ CPU @ 2.80GHz processor and 32 GB RAM. The reported results are an average of three measurements for each test.

## Results

DiviSSR takes a novel approach to identify perfect DNA tandem repeats. The algorithm relies on numerical transformation of DNA sequences and application of simple arithmetic. DiviSSR’s implementation of basic arithmetic is computationally inexpensive when compared to string comparison, regular expression or combinatorial approaches employed by other tools. Our algorithm is 100% exhaustive and identifies all instances of tandem repeats within the user-defined criteria. The time complexity of DiviSSR is *O(nm)*, where *n* is the input data size and *m* is the number of desired motif sizes. We employ two tests for comparison of run time profiles based on these two parameters across tools.

### Comparison of run times

We first checked the effect of input genome size on the performance of DiviSSR and other tools. We selected 5 genomes of different genome sizes as shown in Table 1. Fig 2(i) is a set of bar plots showing the times taken by all tools for organisms of different genome sizes. For all tools, the time taken is linearly dependent on the genome size, but the rate of increase is different. DiviSSR has the least run times in all cases. The speed gain is mainly because DiviSSR uses basic arithmetic and bitwise operations where possible. The slope of the increase in time taken against genome size is also lower for DiviSSR in comparison with other tools. This indicates that our algorithm is highly scalable for large genomes and is even suited for identifying repeats in large sequence datasets.

**Table 1:** Genomes selected for run time tests

| Organism | Common Name | Genome Size (Mbp) |
| --- | --- | --- |
| *S. cerevisiae* | Yeast | 12.3 |
| *C. elegans* | Worm | 102.29 |
| *G. gallus* | Chicken | 1086.67 |
| *D. rerio* | Zebrafish | 1712.79 |
| *H.sapiens* | Human | 3150.05 |

**Figure 2:**
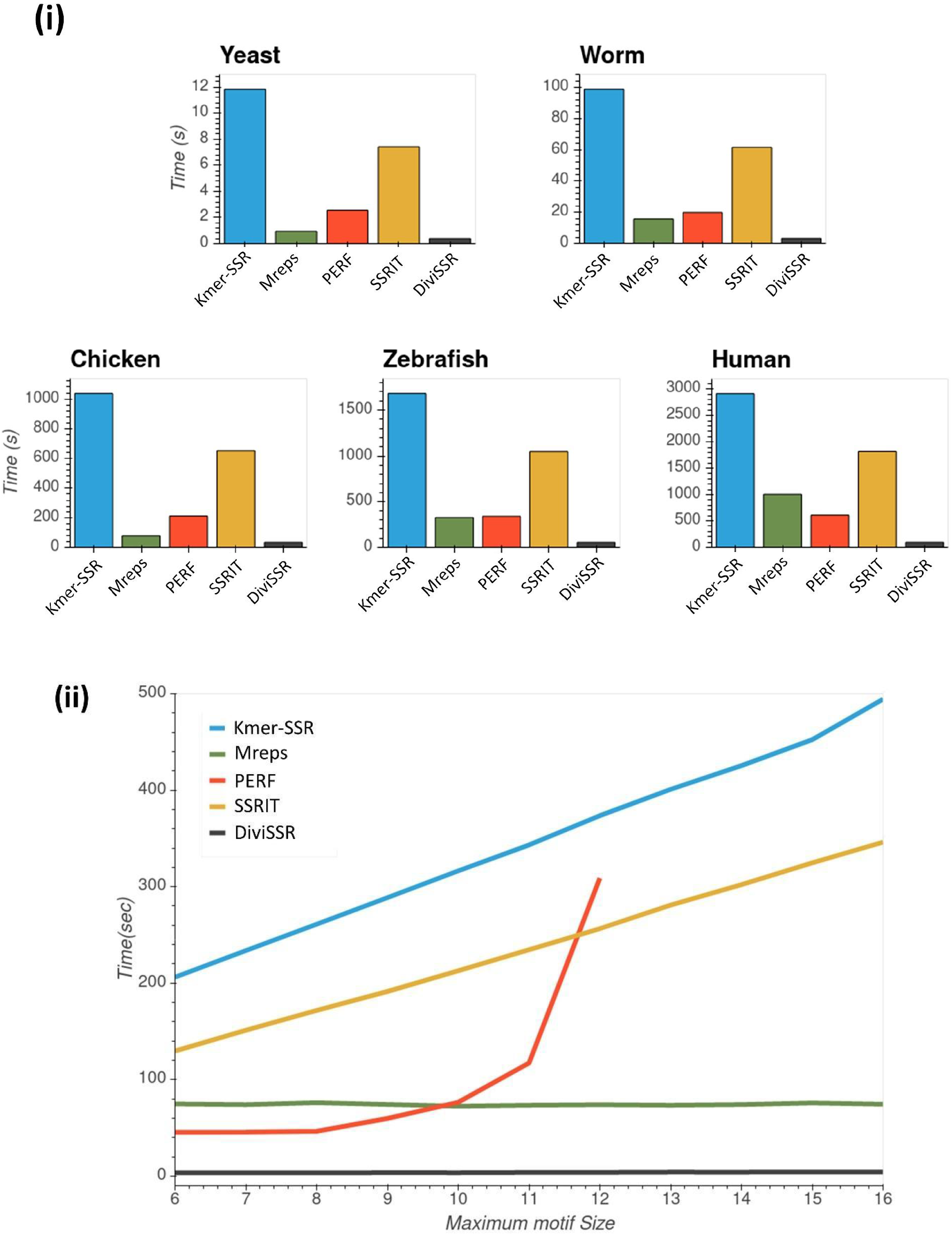
(i) Effect of genome size on run times of different tools. X-axis: Genome size ; Y-axis: Time taken in seconds. (ii) Effect of maximum desired motif size on run time of various tools. X-axis: Maximum motif length (6bp - 16bp); Y-axis: Time taken in seconds.

We next compared the tools’ performance based on the query motif sizes. This was carried out by changing the identification parameters, i.e. desired motif size range of the repeats across all tools. The aim was to analyse the variation of runtime for DiviSSR based on the second parameter it is dependent on. The performance of each tool was profiled on human chromosome 1 (hg38 build) with motif size ranges of 1-6bp, 1-7bp, and so on upto 1-16bp. DiviSSR avoids redundant checks based on the factor-multiple relationship between the desired motif sizes as explained in Materials and Methods. Therefore, the run time of DiviSSR remains relatively constant even as the number of desired motif sizes increases. In comparison with other tools, DiviSSR’s run time remains the lowest at all size ranges (Fig 2(ii)). It is interesting to note that Mreps shows a constant time as the motif size range increases, also making it scalable for longer motif sizes.

### Genomic Annotation

Several studies have demonstrated the gene regulatory functions of SSRs, and a recent report correlates the repeat length to expression of proximal genes [13]. Thus, understanding the genomic context of the repeat is as important as identifying it. DiviSSR is packaged with an auxiliary function which annotates each repeat with its genomic context when provided with the feature information. DiviSSR accepts both GFF and GTF formats as an input for genomic feature information. Downstream to identification of repeats, DiviSSR annotates each repeat with the nearest gene. Based on the overlap of the repeat within a gene it is annotated in the preferential order as either “Exon’’ or “Intron”. Repeats with no overlap with a gene are reported as “Intergenic”. In addition, repeats are also annotated if they occur in the promoter regions of the gene. The promoter region for a gene can be user defined, the default being 1000bp upstream and downstream of the TSS. The genomic annotation for each repeat is denoted in seven additional columns in the output, as explained in Supplementary information.

### Compound repeats

Compound repeats are two or more tandem repeats that either overlap with each other or occur at a close distance to each other. DiviSSR has an additional feature to identify compound repeats with customisable parameters for distance between individual repeats. The output for compound repeats follows a columnar format similar to that of simple repeats but is reported in a separate file. The output format is chosen to be machine friendly and easily parsable using simple regular expressions (See supplementary). When the option of compound repeats is chosen, DiviSSR merges repeats as it scans through the input sequence by storing the location of the previous repeat. This optional merging function has a negligible effect on the total runtime (Fig S2).

### Repeats in raw sequencing datasets

DiviSSR is one of the first tools that can analyze and retrieve repeats from raw sequencing datasets. We have tested the performance of DiviSSR on three test input raw sequencing datasets (Table 2). Though the overall sequence content was similar in all the three datasets (7.5 Gbp), the rate of processing was not proportional. This could be because of the additional overhead cost of processing more reads in datasets of smaller read lengths. DiviSSR processes a typical 30X human genome data (∼100 Gbases) in ∼30min (data not shown).

**Table 2:**
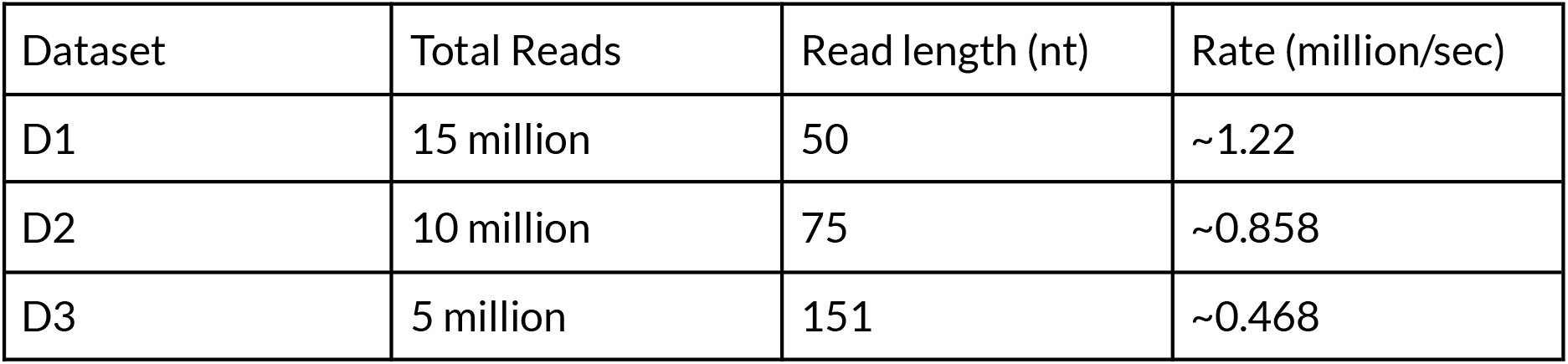
Details of the raw sequencing datasets used to test DiviSSR.

### Format specific reporting

The reporting differs between both the input formats. In case of FASTA input, DiviSSR reports the chromosomal location for each identified repeat following the widely used BED format. Besides the location, the motif sequences, length of the repeat and other characteristics are mentioned in additional columns. For FASTQ input, the location of each repeat is meaningless in the context of individual reads. Hence, DiviSSR reports an overview of repeat composition and repeat type specific statistics for a raw sequencing dataset. Overview statistics include total number of repeat instances, total number of reads with repeats. Similar statistics specific to each repeat type are reported in tabular format, with each row representing statistics for a different repeat motif. The output format for both the input types is explained in detail in the supplementary information.

### Analysis report

To facilitate easy downstream analysis of the output, DiviSSR creates an interactive HTML report summarising the repeat distribution in the input sequence data. The layout of the interactive report is similar for both the input formats though the reported information differs slightly. The report is divided into four sections. The first section has basic sequence information of the input file. The genome size, number of chromosomes, genomic GC content for a fasta file. While for a fastq file the number of reads, the total number of bases and a histogram depicting read length distribution are displayed. The second section summarises the repeat statistics, such as total number of SSR instances, number of bases covered by SSRs, and the normalised frequency and density of SSRs. The SSR stats are normalised by the total number of bases in both fasta/fastq formats for fastq the stats are additionally normalised by the total number of reads. The following section displays a table with repeat class specific summary statistics such as frequency and base abundance. For fasta input format, this section contains two additional tables with the details of top 100 longest repeat instances and top 100 repeat instances with most units. The final section is identical for both input formats and has three switchable tabs each one displaying an interactive chart. The plots summarise the distribution trends of each repeat class based on characteristics such as type of the repeat and length. The detailed explanation of each plot is provided in the supplementary information.

## Discussion

Identification of simple sequence repeats has been solved many times before by different approaches with improvements in performance. Established repeat identification algorithms rely on computationally heavy string-matching algorithms or implement complex data structures. Thus, the solutions so far have not been scalable for analysing large genomes or sequencing datasets. Our present work describes a method which identifies tandem repeats by applying basic arithmetic operations on transformed DNA sequences. The 2-bit representation brings down the number of required operations to be performed on a sequence by four-fold, and also allows application of native bitwise operations for sequence processing. Our implementation of this algorithm can process the whole human genome in ∼30 secs, which is five times faster than the previously reported best [16]. To the best of our knowledge, DiviSSR is also the first tool to identify repeat sequences in raw sequencing datasets and filter the reads with repeat sequences.

Numerical transformation of nucleotide sequences and application of established mathematical methods has been used for predictions of coding sequences, DNA sequence structures and estimating sequence similarity. While most of these methods rely on predictive mathematical models, the backbone of DiviSSR’s algorithm is a computationally inexpensive arithmetic operation, divisibility check. We foresee that similar methods can be used in efficient identification of other DNA elements such as inverted repeats, adapter sequences, restriction sites/palindromes. We believe these tactics can even be applied to classic computational biology problems, such as sequence alignment, and improve processing time bottlenecks in other genomics pipelines.

## Acknowledgements

The authors thank Prof Rahul Siddharthan for critical reading of the manuscript and providing valuable suggestions. AA is a recipient of a Senior Research Fellowship from the Council of Scientific and Industrial Research.

