## Supplementary for "DiviSSR: Simple arithmetic for efficient identification of tandem repeats"

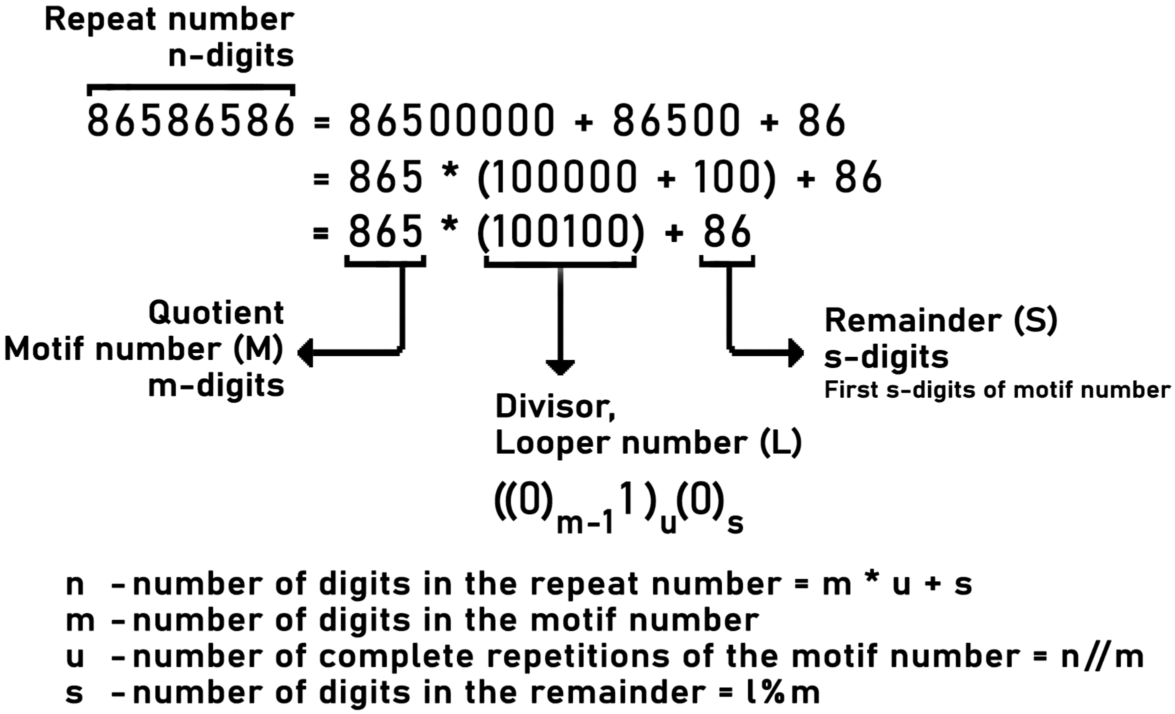


**Figure S1**: Breaking down a repeat number (86586586) into unit parts to illustrate the generalised numerical property. The number 86586586 is written as 865*100100 + 86 where 865 becomes the motif number and 100100 becomes the repeat base or the DiviSSR.

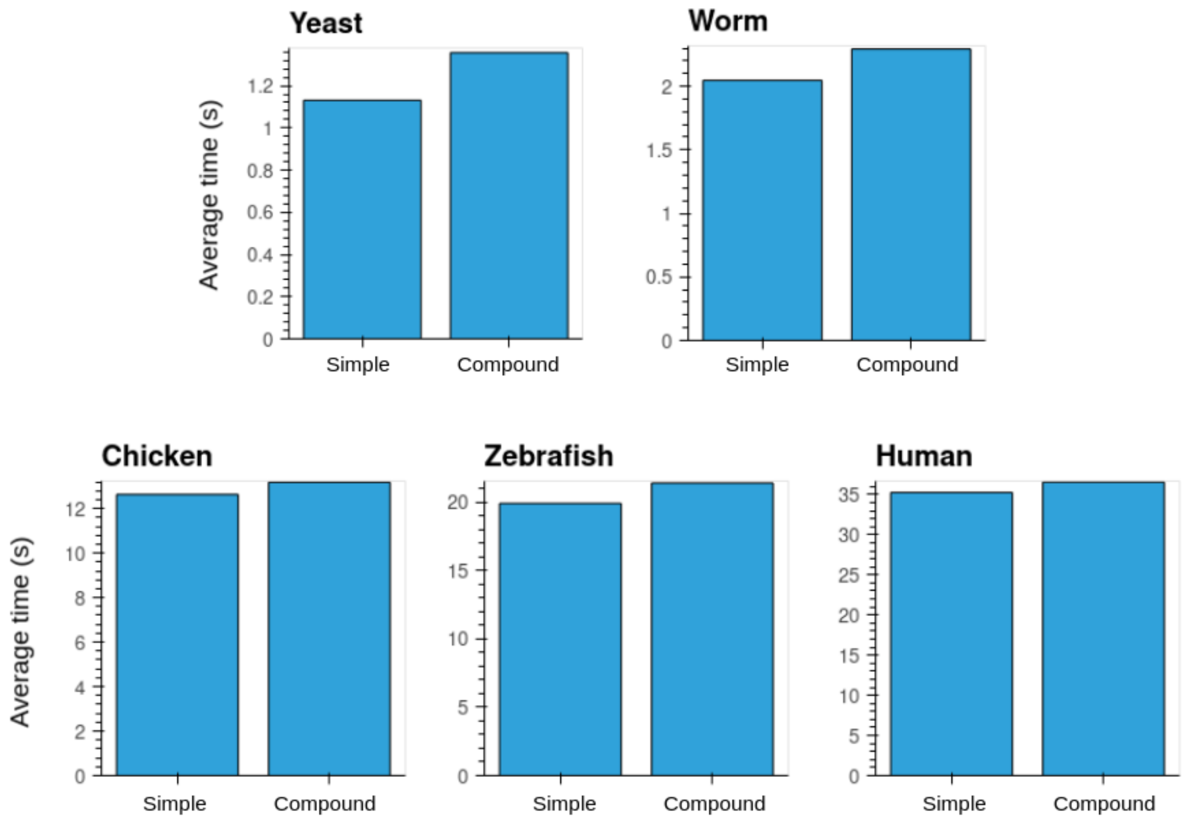


**Figure S2:** Difference in average time taken by diviSSR to identify simple repeats and compound repeats.

### *Repeat class*

Repeat class is used for categorisation of repeat sequences based on the motif sequence. A repeat class represents all repeats of motifs which are cyclical variations of itself or cyclical variations of the repeat class. For example repeats of the motifs ATG, TGA, GAT, CAT, ATC, and TCA are all categorised belonging to the repeat class ATC. The repeat class is chosen lexicographically i.e., the alphabetically first motif is selected as the repeat class. In our 2-bit conversion we also assign the numbers to nucleotides lexicographically i.e., A=00, C=01, G=10, and T=11. This retains that motif which is first lexicographically has the lowest numerical value. To get the repeat class of a motif we generate all its cyclical variation and cyclical variations of reverse complements and check for the lowest number in the 2-bit representation. We save the repeat class for a motif as a map to avoid calculating the repeat class everytime we encounter a repeat of that motif.

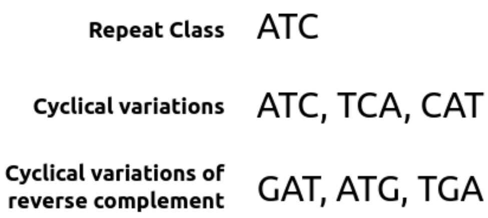


**Figure S3:** Categorisation of motifs into a repeat class based on cyclical relationship.

### *Algorithm for repeat annotation*

Here we explain the algorithm employed for genomic annotation of repeats post identification. The input genomic feature file can be both GFF/GTF. DiviSSR processes the feature file and builds two objects, one with chromosome wise gene information and the other with gene wise exon information. The first object stores the list of all genes sorted in ascending order of the start position for each chromosome. For each gene its start and stop positions, strand orientation and a unique gene identifier are stored. The subgene object stores a map of all the exon positions in ascending order for each gene against the gene's unique identifier. As DiviSSR reports repeat locations in a position based sorted order, sorted storage of gene information allows unidirectional check of possible annotation.

The possible annotations of repeat are depicted in the Fig S3. The aim is to either find an overlapping gene or the closest non-overlapping gene for each repeat. For a gene with start position *GS* and end position *GE*, and a repeat with start position *RS* and end position *RE*, the genic/promoter overlap is possible only for genes which have either $GS\text{-}P < RS < GE+P$ or $GS\text{-}P < RE < GE+P$. The unidirectional search for the desirable gene optimises the search as a gene not overlapping with the current repeat will not overlap with following repeats. When we start with the first repeat on the chromosome from the list of genes on the chromosomes we start comparing the repeat position with the position of each gene. For every gene where the repeat goes beyond the gene is ignored and the repeat is annotated with respect to the gene for which the overlap is found. If the overlap is not found the repeat is annotated as distal intergenic with the closest gene. For this comparison we save the index (*GI* ) of the latest gene on the chromosome where the repeat falls beyond the allowed range. The following repeat is checked for overlap only with genes from and beyond the index *GI* on the chromosome. The priority of annotation is for a repeat is given for Promoter, Exonic, Intronic and Distal Intergenic.

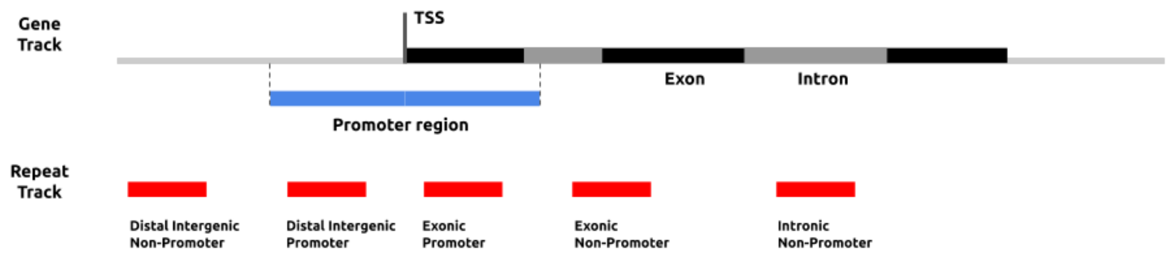


**Figure S4:** Shows the annotation tag of each repeat based on the position w.r.t the gene and its sub features.

### *Bitwise operations for sequence representation after shifting window*

As the window is shifted by 1bp at each step, concurrently the binary number representing the DNA sequence undergoes 3 bit-wise operations to reflect the shifted sequence. First it is shifted by 2-bits to the left, second the binary representation of new nucleotides is added by an OR operation. At this point this number is a representation of $k$+1 nucleotides. To represent only the latest $k$ bases we perform an AND operation with the binary number $2^k$-1 and retain the last 2$k$ bits.

### *Format specific reporting*

The difference in output format for based on the input sequence format was mentioned in the results section. The following tables delineate the column wise information reported for both the formats.

| *FASTA FORMAT* | |
|---|---|
| **S.no** | **Description** |
| 1 | Sequence identifier |
| 2 | Start position of the repeat |
| 3 | End position of the repeat |
| 4 | Repeat class of the repeat |
| 5 | Length of the repeat in bp |
| 6 | If the actual_repeat is reverse complement of the repeat class |
| 7 | The actual motif of the start of the repeat |
| 8 | Identifier of the gene |
| 9 | Start position of the gene |
| 10 | End position of the gene |
| 11 | Strand orientation of the gene |

| | |
|---|---|
| 12 | Annotation w.r.t the closest gene (Genic, Exon, Intron, Intergenic) |
| 13 | Annotation whether the repeat falls in the promoter region |
| 14 | Distance from TSS of the closest gene |

| ***FASTQ FORMAT*** | |
|---|---|
| **S.no** | **Description** |
| 1 | Repeat class |
| 2 | The total number of instances of the repeat |
| 3 | Number of unique reads with instances of the repeat |
| 4 | Total number of bases covered by the repeat |
| 5 | Total instances of repeats normalised to total number of reads |
| 6 | Reads with repeats normalise total reads in the file |
| 7 | Semicolon separate key=value pairs depicting frequency of repeat at a particular length |

### *Compound repeats output format*

The output report of the compound repeats is similar to the output of simple repeats with changes in the way the repeat sequence is reported. The fourth column which reports the repeat class in the case of reporting a simple repeat, reports the sequential occurrences of individual repeat classes in the compound repeat. A number is associated with each repeat class which denotes the number of consecutive repeats having the same repeat class. For example (AC)1(AGAT)2, the above notation represents a compound repeat which starts with a simple repeat of a motif belonging to class AC, followed by a repeat of a motif of class AGAT and ends with a repeat of a different motif of the same class AGAT. The fifth column reports the total length of the compound repeat. The sixth column reports the strand orientation of the repeat motif with respect to the repeat class. The last column reports the actual motifs, repeat lengths and distances between the individual repeats of the compound repeat in a specific format. Each repeat motif is placed in parentheses followed by the length of the repeat. Two individual repeats are separated by two "|" between which the distance between the two repeats is reported. Example, (AAT)14|D5|(AGAA)23, which represents a compound repeat with an AAT repeat of length 14bp followed by an AGAA repeat of 23bp which are separated by a distance of 5bp. Overlapping repeats are reported with a negative distance between them which represents the number of overlapping bases. This format is chosen to be easily parsable and also regex friendly.

### *Interactive visualisations in HTML report*

The final section in the HTML report is identical for both input formats and has three switchable tabs each one with an interactive chart. The plots are identical to the plots displayed in the analysis report of PERF. The repeat abundance bar chart provides plots of the abundance frequency or bases covered by each repeat class. The repeat distribution is a stacked bar chart, summarising the relative abundance of each repeat based on the motif size. The length versus frequency chart helps to analyse the abundance of each repeat at different lengths of the repeat. These charts provide the user to analyse specific distribution patterns based on the type of the repeat and its characteristics.
